# Rapid Sequencing of Multiple RNA Viruses in their Native Form

**DOI:** 10.1101/482471

**Authors:** Thidathip Wongsurawat, Piroon Jenjaroenpun, Mariah K. Taylor, Jasper Lee, Aline Lavado Tolardo, Jothi Parvathareddy, Sangam Kandel, Taylor D. Wadley, Bualan Kaewnapan, Niracha Athipanyasilp, Andrew Skidmore, Donghoon Chung, Chutikarn Chaimayo, Michael Whitt, Wannee Kantakamalakul, Ruengpung Sutthent, Navin Horthongkham, David W. Ussery, Colleen B. Jonsson, Intawat Nookaew

## Abstract

Long-read nanopore sequencing by a MinION device offers the unique possibility to directly sequence native RNA. We combined an enzymatic poly-A tailing reaction with the native RNA sequencing to (i) sequence complex population of single-stranded (ss)RNA viruses in parallel, (ii) detect genome, subgenomic mRNA/mRNA simultaneously, (iii) detect a complex transcriptomic architecture without the need for assembly, (iv) enable real-time detection. Using this protocol, positive-ssRNA, negative-ssRNA, with/without a poly(A)-tail, segmented/non-segmented genomes were mixed and sequenced in parallel. Mapping of the generated sequences on the reference genomes showed 100% length recovery with up to 97 % identity. This work provides a proof of principle and the validity of this strategy, opening up a wide range of applications to study RNA viruses.

## INTRODUCTION

Infectious disease epidemics are primarily driven by RNA viruses (Woolhouse et al., 2016; Carrasco-Hernandez et al., 2017) and hence are likely agent of future pandemics (Carrasco-Hernandez et al., 2017). Due to the lack of proofreading in RNA polymerases, RNA viruses are error-prone during genome replication, providing a platform for rapid evolution in new environments and hosts. Currently, genome sequences of many viruses are available and served as a powerful resource for molecular surveillance, pathogen characterization, diagnosis, and antiviral drug discovery (DeFilippis et al., 2003). Still, outbreaks of viral diseases are a never-ending challenge. Short-read sequencing methods to recover RNA viral genomes relies on reverse transcription (RT) of RNA to cDNA which requires primer optimization and amplification. These steps introduce many biases, artifacts and makes rapid diagnosis difficult (Marston et al., 2013). Alternatively, real-time sequencing using a pocket-sized MinION device Oxford Nanopore Technologies (ONT) skips these steps and makes rapid detection and characterization of emerging pathogens possible (Hoenen et al., 2016; Kilianski et al., 2016; Faria et al., 2017). A shorter procedure on the MinION by elimination of cDNA synthesis step can be accomplished through direct RNA sequencing (dRNA-seq), designed to directly ligate and sequence only polyadenylated RNA, providing strand-specific information and base modifications of RNA (Garalde et al., 2018; Jenjaroenpun et al., 2018). Independent from the poly(A) enrichment, a sequence-specific 3’-adapter for targeting non-polyadenylated RNA can be designed and has been successfully demonstrated for rRNAs (Garalde et al., 2018) and influenza virus (Matthew W Keller, 2018). This method can reduce background noise from host RNAs. However, designing sequence-specific 3’-adaptor needs a prior knowledge of a nucleotide on that end of the target as prerequisite. Therefore, undiscovered RNA viruses, or coinfections might not be discovered by this approach.

We established a rapid procedure and evaluated the feasibility of MinION for sequencing of multiple, single-stranded RNA (ssRNA) viruses in a single reaction to obtain complete genomes. Our approach combines (1) poly(A)-tailing using simple enzymatic reaction, (2) native RNA sequencing by MinION, and (3) real-time analysis aim to simultaneously detect and characterize multiple RNA viruses. A mix of negative(-) and positive(+) ssRNA viruses, including Mayaro virus (MAYV), Venezuelan equine encephalitis virus (VEEV), Chikungunya virus (CHIKV), Zika virus (ZIKV), Vesicular stomatitis virus (VSV), and Oropouche virus (OROV) were used in this demonstration.

## RESULTS

To establish an optimal procedure for real-time native RNA sequencing, we performed three experimental scenarios. 1) Sequencing two polyadenylated RNA viruses without RT, suggested by the original protocol, 2) Adding poly(A)-tail to non-polyadenylated RNA phage and ZIKVs using *E. coli* poly(A) polymerase (EPAP), followed by sequencing, and 3) Applying ribosomal RNA (rRNA) depletion step to host-rRNA contaminated samples, followed by poly(A)-tailing and sequencing. Then, we applied the optimized protocol on a mix of RNA viruses.

In the first scenario, naturally, polyadenylated (+)ssRNA alphaviruses, MAYV and VEEV, genome size approximately 11.5kb, were used. Extracted RNA (Materials and methods) of MAYV and VEEV were pooled at approximately the same concentration and used for library preparation of the dRNA-seq protocol, following the manufacturer’s recommendations, except the RT step was skipped (Materials and methods). The prepared library then was loaded into ONT flow cell and sequenced. The sequenced reads were mapped on the genomes and analyzed (Table 1), resulting in full genome coverage of the both viruses with 716X and 8184X, for MAYV and VEEV, respectively. We obtained 97% sequence identity by consensus polishing. A high viral/host ratio of 21 was observed because the most abundant RNA species in host, rRNAs, were not captured. By this scenario, multiple viral whole genome sequencing of polyadenylated viruses without RT is possible.

**Table 1.** Three optimization scenario.

| Scenario | Virus | Genome size (kb) | RNA input (ng in total) | Coverage | Mean coverage | Longest mapped length (kb) | Mapped virus (%) | Mapped host, % (host rRNA) | Low quality sequence (%) | virus/host (fold) | Total number of reads |
| --- | --- | --- | --- | --- | --- | --- | --- | --- | --- | --- | --- |
| Multiple viral detection | MAYV<sup>a</sup> | 11.14 | 350 | 100% | 716x | 11.13 | 6.35 | 4.13 (0.41) | 8.45 | 21.1740 | 92688 |
|  | VEEV<sup>a</sup> | 11.45 | 350 | 100% | 8184x | 11.38 | 81.07 |  |  |  |  |
| An enzymatic reaction for non-poly(A)-tail viral RNA | phage MS2 | 3.57 | 500 | 100% | 2887x | 3.54 | 71.58 | 0 (0) | 28.42 | NA | 9553 |
|  | ZIKV | 10.79 | 54 | 100% | 4x | 10.32 | 0.11 | 81.92 (71.32) | 17.97 | 0.0014 | 16793 |
| Host rRNA depletion and enzymatic reaction for non-poly(A)-tail viral RNA | ZIKV | 10.79 | 48 | 100% | 12x | 10.53 | 3.98 | 17.59 (3.32) | 78.43 | 0.2264 | 904 |
|  | 2% phage MS2 + 98% mouse total RNA | 3.57 | 60 (phage), 2940 (mouse) | 100% | 426x | 3.47 | 3.78 | 34.93 (0.35) | 61.29 | 0.1083 | 32984 |

MAYV, Mayaro virus; VEEV, Venezuelan equine encephalitis virus; ZIKV, Zika virus
<sup>a</sup>Presence of polyadenylated genome

The second scenario sought to evaluate the feasibility of using the poly(A)-tailing reaction coupled with dRNA-seq, (+)ssRNA. Bacteriophage MS2 was used first. Five hundred nanograms of 3.57kb RNA was treated with EPAP using an optimized poly(A)-tailing reaction (Materials and methods). Poly(A)-tailed RNA was then used as input for library preparation and sequencing in the same way as described above, resulting in a complete read of the bacteriophage genome (Table 1). We then applied the procedure on ZIKV RNA, non-polyadenylated (+)ssRNA with ~11kb length, and obtained long-read. In both scenarios, we observed reduced coverage at the 5’ and a heavy coverage bias towards the 3’ however we found reads that almost spanned the entire genome (Figure 1B). By poly(A)-tailing reaction, the majority of RNA in ZIKV sample was host RNA that could be captured after EPAP treatment resulted very low virus/host ratio of 0.0014. By this scenario, we obtained a rapid procedure to universally detect ssRNA viruses in real-time. The workflow of poly(A)-tailing plus dRNA-seq was accomplished and summarized in Figure 1A. The host RNA contamination problem was taken into account in the next scenario.

**Figure 1.**
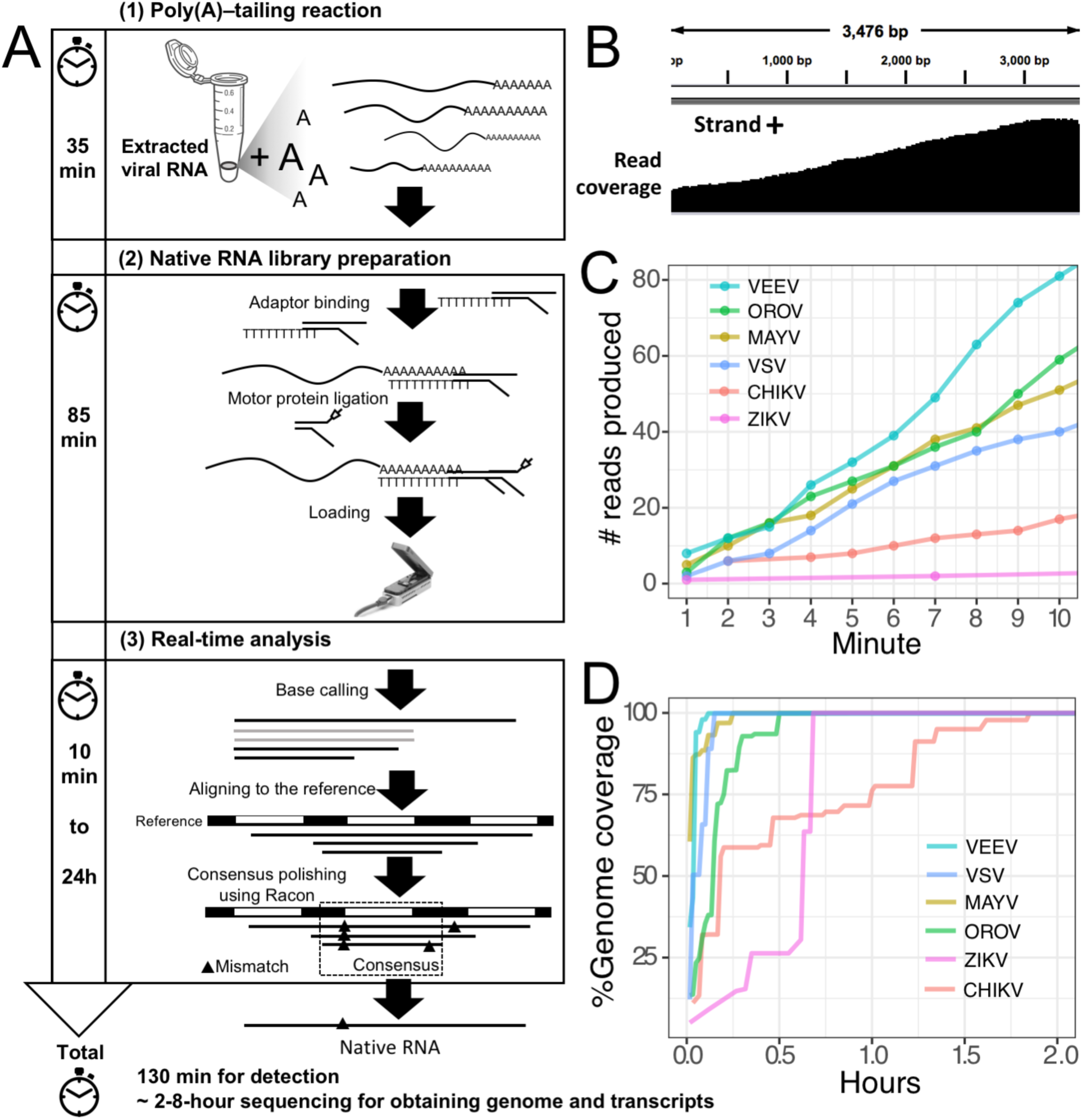
Poly(A)-tailing native RNA-seq protocol. (A) The protocol contains three main steps 1) Poly(A)-tailing reaction using *E.coli* poly(A) polymerase (EPAP) 2) native RNA-seq by nanopore MinION and 3) real-time analysis to retrieve native RNA sequence. (B) An example of sequencing result (reads) that mapped to the bacteriophage MS2 genome. (C) For detection purpose, this protocol can provide mapped RNA sequences within the first two minutes after sequencing. (D) To obtain whole genome and mRNAs, 2-8 hours sequencing could provide more complete information.

In the third scenario, we introduced host rRNA depletion step using a rRNA removal kit, to increase the amount of viral RNA reads. The rRNA depletion was performed before poly(A)-tailing and dRNA-seq. The ZIKV sample was used to evaluate changes in ratio of viral/host RNA. We found that introducing the depletion step resulted in a 160-fold increase in proportion of viral RNA reads/ host reads compared to the non-depleted sequencing. However, we observed high proportion of low-quality sequences compared to non-depleted scenario. We further tested the impact on rRNA depletion step on artificial mixing of 2% of bacteriophage MS2 RNA and 98% of total RNA extracted from mouse cells. The rRNA depletion kit was used for the mixture as described above. Again, the proportion of low quality reads was increased as seen in rRNA-depleted ZIKV sample (Table 1). This problem may be come from the remained compound from the rRNA depletion process that interfere the dRNA-seq chemistry. Therefore, the depletion step need to be further optimized, we decided not to use the depletion step in our protocol, as it may add complexity and increase the low-quality sequences.

Finally, we applied the established procedure to sequence a pool of six RNA viruses including MAYV, VEEV, CHIKV, ZIKV, VSV, and OROV, which were each purified from supernatant of varying cell lines and RNA was extracted from purified virions (Materials and methods). This mixed RNA viruses has diversified genomic property i.e., (+)ssRNA, (-)ssRNA, lack/presence of poly(A)-tail, segmented/non-segmented genomes (Table 2). The RNA input from each sample varied from 16 to 66 ng (215 ng in total) (Table 2). Sequencing on MinION was performed for ~8 hours in total with real-time base calling, enabling data analysis in real-time (Materials and methods). The sequences were mapped to the viral genomes and host genomes (Table 2). Remarkably, within the first two minutes, the first reads of all six viruses were translocated through nanopore (Figure 1C). In only two hours, complete or near complete genomes of all the six viruses were acquired (Figure 1D). In total 85,711 reads were generated, 5,000 (6%) were mapped on viruses, 62,060 (72%) were mapped on host genomes, and 18,651 (22%) were unclassified (low quality). Mapping of the 5,000 reads on the six genomes showed 100% recovery of the MAYV, VEEV, CHIKV, VSV, and OROV at 48X – 659X coverage and showed 99% recovery of the ZIKV, the least abundance in the mix, at 10X coverage (Table 2). We observed several long reads (>10kb) that covered > 95% of MAYV, VEEV, and VSV genomes while the longest reads mapped to CHIKV and ZIKV were 84% and 68% of genome length, respectively. Unlike the other viruses, OROV has a tripartite genome of small (S), medium (M), and large (L) segments. We found the longest read covered 99% of S (0.96kb), 98% of M and L (4.39kb and 6.85kb). Notably, although high error rates of individual native RNA sequence were observed (~82% read identity), 93–97% identity can be achieved by consensus polishing.

**Table 2.**
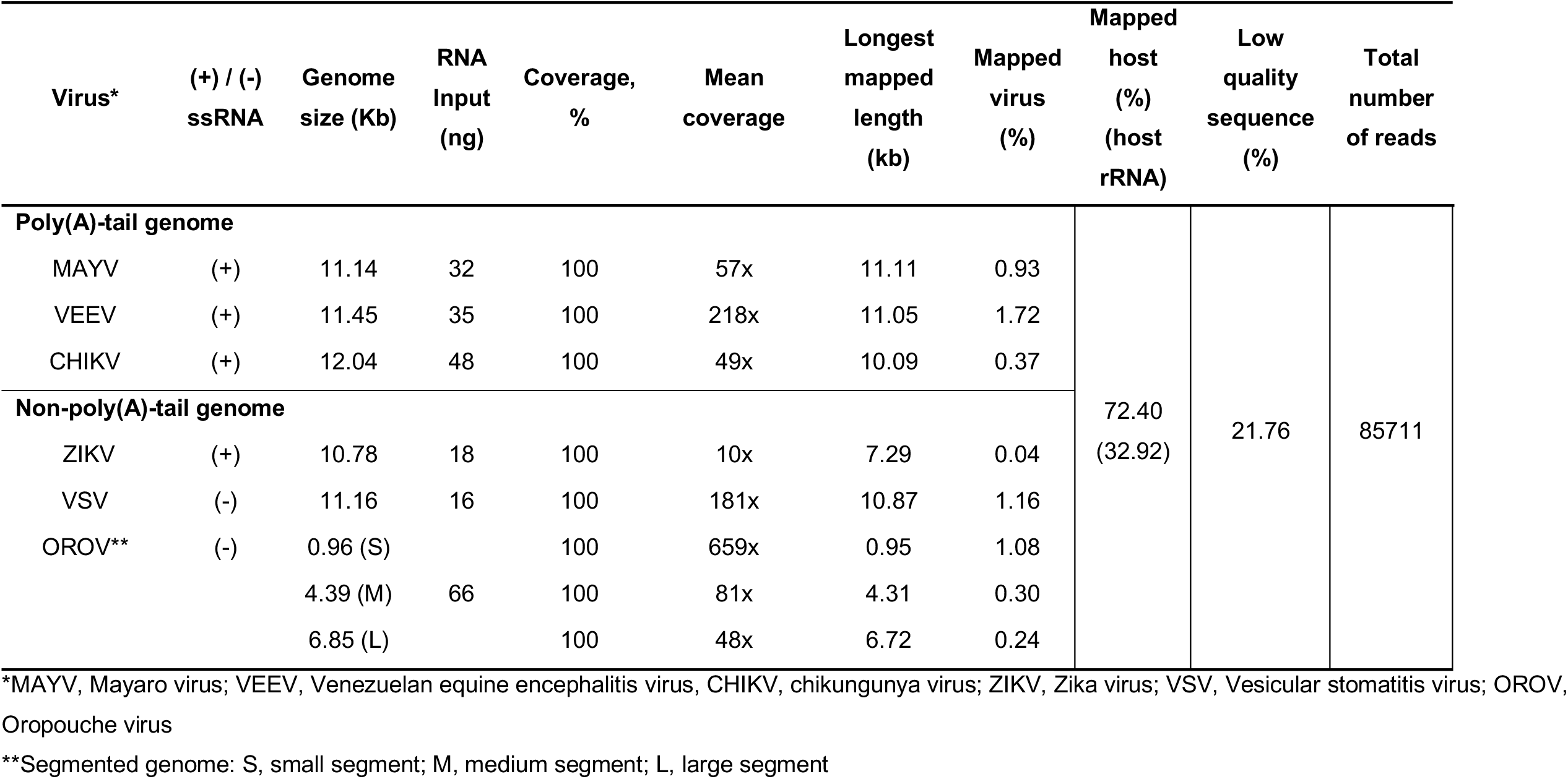
List of virus used in this study and sequencing result.

We inspected the mapped reads based on strand-orientation and found feasibility of simultaneous detection of viral genome and its transcripts (Figure 2). We observed the two bumps around the gap of two subgenomic mRNA of VEEV on the coverage plot (Figure 2A) clearly indicating that the two subgenomic mRNAs were independently transcribed and the reads that spanned over the gap were belong to the genomic material (Materials and methods). This result based on sequencing RNA directly without primer bias, reverse transcription and amplification. However, observation of two subgenomic mRNA in VEEV need further deep investigation. Conversely, (-)ssRNA virus needs to transcribe mRNA in the opposite direction of the genome. We found mapped reads derived from the mRNA and genome of OROV nicely aligned in opposite direction (Figure 2B).

**Figure 2.**
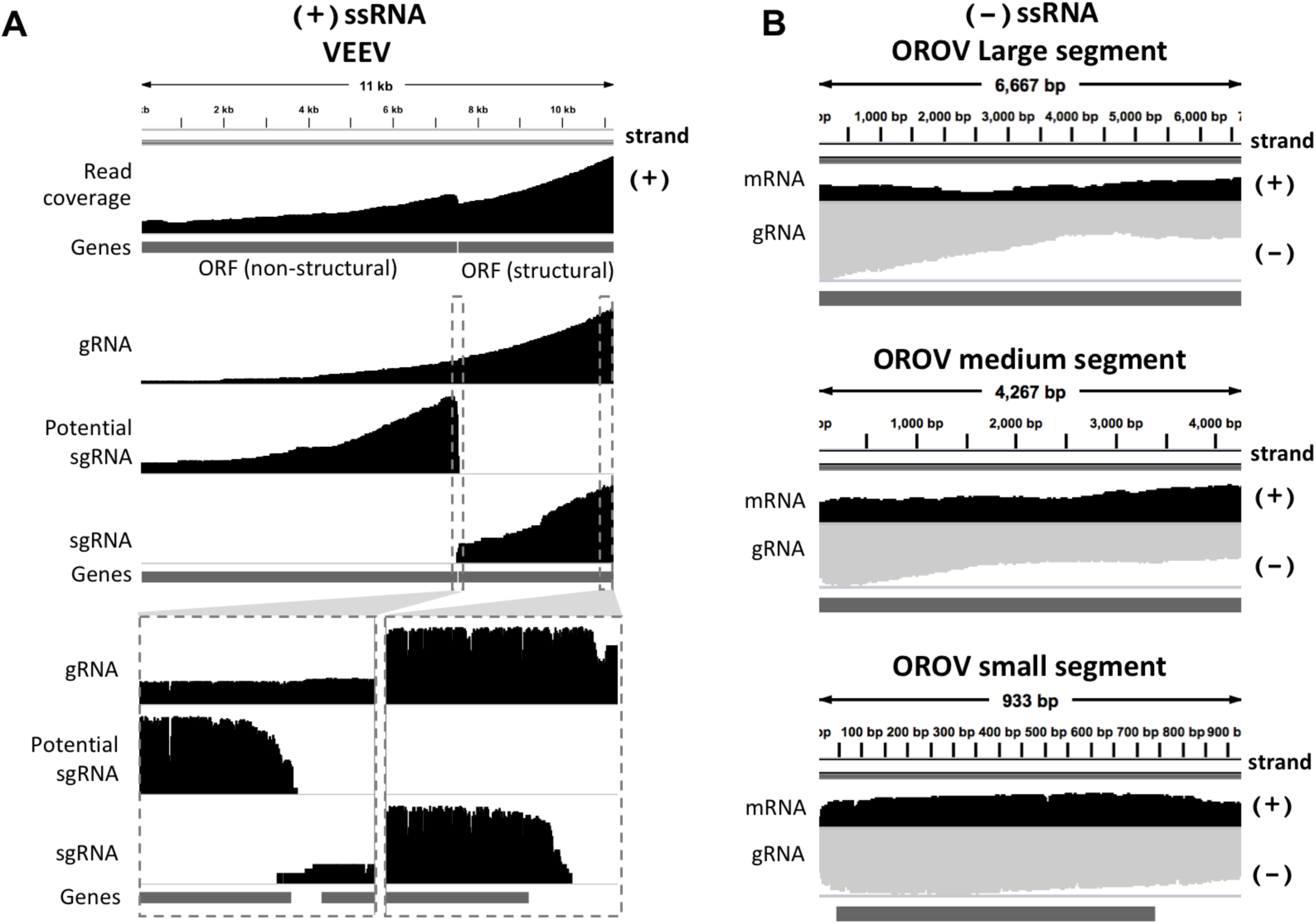
Strand-specific sequencing of (+)/(-)ssRNA. (A) Mapped sequences (coverage depth) of (+)ssRNA genome VEEV and subgenomic RNA were plotted in the same direction as shown integrative genomics viewer (IGV). The top panel shows all mapped reads. The middle panel shows the three category of mapped reads belong to genome (gRNA) and the two subgenomic mRNA(sgRNA). The bottom panel shows the zoom in view of non-subgenomic mRNA region that enable discrimination between genome and subgenoimic RNA (B) Mapped sequences (coverage depth) of (-) ssRNA OROV genome (gRNA) and mRNA were plotted in the opposite direction. Black: positive strand and Grey: negative strand. The dark grey boxes in the bottom of each panel were gene annotation. Top, middle and bottom panel illustrated the three segment S, M and L of OROV, respectively.

## DISCUSSION

This versatile protocol can contribute viral RNA research by (i) retaining the strand direction; this facilitates observation of genome, subgenomic mRNA/mRNA, recombinant, complementary RNA intermediates (ii) increasing the accuracy of determining transcriptomic architecture and recombination with no need for assembly (iii) providing a fast and easy method to perform direct sequencing of any RNA virus (iv) directly sequencing of novel RNA viruses with no specific primer required, and (v) saving on the budget (see Table 3). In addition, the native RNA signal can be used for further detection of RNA base modifications that are known to control viral replication during host infection (Lichinchi et al., 2016). The ability to generate multiple levels of genetic information at high speed has potential to increase our basic understanding of the known and undiscovered viruses at a rapid rate. Although promising, however, this protocol carries a number of limitations. First, this protocol requires high amounts of virus. Second, the poly(A)-tail reaction is not only specific to viral RNA; host RNA can also be a target. Competition between host RNAs and viral genome requires a better strategy for host RNAs depletion. Third, although we could identify virus at species-level from error-prone long reads, deep sequencing is required for accurate strain-level identification. However, improvement of nanopore read accuracy is subject to extensive research, and it is likely that much better accuracy will be available in the future, as well as better depth of viral reads, leading to strain-level identification. In conclusion our simple protocol could provide rapid viral RNA genome/subgenome sequencing and characterization that can be a versatile tool to study biology of RNA viruses.

**Table 3.** Advantages and limitations of this protocol

| **This protocol enables to:** | **Advantages** | **Limitations/ difficulties** |
| --- | --- | --- |
| (i) detect complex population of RNA viruses simultaneously whether (+)ssRNA, (-)ssRNA, lack/presence of poly(A)-tail, segmented/non-segmented genomes | - No specific primer design<br>- Co-infection/co-circulation detection<br>- Save on the budget | - Require high viral load from cultivation<br>- Poly(A)-tail reaction is not only specific to viral RNA; host RNA can be a target |
| (ii) full-length genomes, subgenomic(sg) mRNAs and mRNAs | - Virus identification<br>- Map transcript architecture and recombination without the need for assembly | - Because of high error rate, strain level detection is still limited<br>- Fully intact RNA and gentle handling are required |
| (iii) fast and easy | - Eliminate primer design, reverse transcription, and PCR bias enabling real-time detection within 3 hours | - A loop structure forms at the 3' terminus might create difficulty for the enzymatic reaction |
| (iv) strand-specific RNA-seq | - Retain the information pertaining to strand origin; facilitate observation of genome/ transcript/ complementary RNA strand intermediate. |  |
| (v) sequence native RNA* | - Provide signal of RNA base modifications | - Only some known modifications are found<br>- Deep sequencing is required for detection of RNA modifications and structures |

\*Possible for study RNA epigenetics in the future.

## Data availability

The sequence native RNA data of six viruses has been submitted to GenBank and is publicly available under the Bioproject number PRJNA493665.

## AUTHOR CONTRIBUTIONS

I.N. and T.W. designed and conceived the project. I.N., C.J. and D.W.U. supervised the study. C.J., M.T., J.L., A.T., J.P., B.K., N.A., A.S., D.C., C.C., M.W., W.K., R.S., N.H. performed viral cultivation and RNA extraction. T.D.W. and T.W. optimized library preparation protocol. T.W. and S.K. conducted MinION sequencing. P.J. and I.N. performed computational analysis. All authors participated in interpreting the results and writing the manuscript, and read and approved the final version of this manuscript.

## MATERIALS AND METHODS

### Viral cultivation and RNA extraction

*Mayaro virus (MAYV), Oropouche virus (OROV), and Venezuelan equine encephalitis virus (VEEV)*

MAYV (strain BeAr505411, Genbank no: KP842818.1), OROV (strain BeAn 19991, Genbank no: S: KP052852.1; Segment M: KP052851.1; Segment L: KP052850.1) and VEEV (strain TC-83, Genbank no: L01443.1) were cultivated on Vero cells in MEM with 5%FBS and 1%PenStrep. The infection was made on 80% confluent cells with an MOI of 1 for Mayaro and 0.5 for Oropouche. Both viruses were incubated with cells for 3 days. The viruses were then aliquoted and centrifuged according to protocol. The TRIzol LS extraction (Invitrogen) was carried out following the manufacturer’s instruction for RNA extraction with the exception of adding glycogen after separating the aqueous phase from the organic layer and interphase. Purified RNA was resuspended in 50 µl RNase-free water and stored at −70 °C.

#### Chikungunya virus (CHIKV)

CHIKV (strain 181/25 was obtained from BEI resources, Genbank no: EF452493.1) was propagated in Vero 76 cells (ATCC CRL-1587) grown in MEM-E with 10% FBS. After CPE was monitored, the cell supernatant was harvested and clarified with a low-speed centrifugation at 4,000 x g for 20 min. The cell supernatant was stored at −80 °C until use and viral RNA was isolated from the supernatants. RNA was isolated using RNAzol RT (MRC) with the Direct-zol RNA miniprep kit (Zymo Research) according to manufacturer’s instructions, with one additional wash step.

#### Vesicular stomatitis virus (VSV)

BHK-21 cells were infected at a multiplicity of 0.1 and the culture medium was collected after 22 hours. The culture media were first clarified by centrifuging at 8,000 x g for 10 minutes then the supernatants were layered on top of a 20% sucrose cushion and pelleted at 100,000 x g for 1 hr. The virus pellet was resuspended in Tris-saline, pH 7.4 and stored at −80 °C. The viral pellet was subjected to a Phenol-Chloroform RNA extraction from TRIzol LS (Invitrogen) following manufacturer’s protocol with the exception of adding glycogen after separating the aqueous phase from the organic layer and interphase. Purified RNA was resuspended in 20 µl RNase-free water and stored at −70 °C.

#### Zika virus (ZIKV)

Two strains of ZIKV were used for this study

1. Genomic RNA of ZIKV (strain MR 766, GenBank: AY632535.2) used for method optimization was purchased from ATCC (https://www.atcc.org/products/all/VR-84.aspx).
2. ZIKV (strain PRVABC59 provided by Dr. Barbara Johnson at CDC, GenBank: KU501215) used in pooled viruses sequencing was propagated in Vero 76 cells grown in MEME (GIBCO Laboratories) with 10% fetal bovine serum. After CPE was monitored, the cell supernatant was harvested and clarified with a low-speed centrifugation at 4,000 x g for 20 min. The cell supernatant was stored at −80 °C until use and viral RNA was isolated from the supernatants. RNA was isolated using RNAzol RT (MRC) with the Direct-zol RNA miniprep kit (Zymo Research) according to manufacturer’s instructions, with one additional wash step.

#### Bacteriophage MS2 RNA

RNA from bacteriophage MS2 (Sigma) is composed of 3569 nucleotides (Genbank no: NC_001417.2). MS2 RNA is free of host nucleic acids and proteins (https://www.sigmaaldrich.com/catalog/product/roche/10165948001?lang=en&region=US).

#### Mouse RNA

RNA isolation was performed from a mouse cell line with the Direct-Zol RNA miniprep Kit (Zymo Research) as per manufacturer’s instructions. Total RNA was eluted in 50µl nuclease free water and stored at −80°C.

### Ribosomal RNA (rRNA) Depletion

Host cytoplasmic and mitochondrial rRNA was depleted from the samples using Ribo-Zero Gold kit (Illumina) according to manufacturer’s instructions. Briefly, magnetic beads were washed, samples were treated with rRNA removal solution, rRNA was removed, and rRNA-depleted samples were purified.

### RNA quantification

RNA was quantified using a Qubit ^®^ 2.0 Fluorometer (ThermoFisher Scientific) and Nanodrop Spectrophotometer (ThermoFisher Scientific). The Qubit^®^ assay uses a Qubit RNA BR (Broad-Range) Assay Kit (ThermoFisher Scientific).

### Poly(A)-tailing of RNA using *E. coli* Poly(A) Polymerase (EPAP)

To obtain high quality RNA for further preparation, extracted RNA of each virus received from a different independent lab was purified and concentrated using AMPureXP beads (Beckman Coulter) and eluted in 15 µl nuclease-free water (NEB). Purified and concentrated RNA was incubated at 70 °C for 2 min and immediately placed on ice to avoid secondary structure formation. Poly(A)-tailing reaction using *E. coli* EPAP (NEB# M0276) was designed to perform in a 26 µl reaction (reaction size can be scaled up as needed). Purified RNA was treated with 2.5 µl 10X *E. coli* Poly(A) Polymerase Reaction Buffer (supplied with the enzyme), 4.5 µl ATP (10mM) (supplied with the enzyme), and 4 µl EPAP (NEB) at 37 °C for 10 min. Poly(A)-tailed RNA was then cleaned using 2.5x (by volume) AMPureXP beads (Beckman Coulter) and eluted with 11-15 µl nuclease free water (NEB), 2 min at room temperature. Cleaned Poly(A)-tailed RNA was incubated for 1 min at 70 C and immediately placed on ice to remove secondary structure. The cleaned Poly(A)-tailed RNA was used as input for direct RNA sequencing library preparation.

### Library preparation and direct RNA sequencing by Nanopore

Direct RNA sequencing was performed using the Direct RNA Sequencing Protocol (SQK-RNA001 kit) for the MinION. Some steps of the manufacture protocol were modified. First, the ligation reaction of RNA and adaptor was extended from 10 to 15 minutes. At this step, control RNA (RCS) was not added. Second, the reverse transcription (50 minutes) step was skipped in our protocol. Third, wire-bored tips (Axygen) were used for mixing AMPureXP beads (Beckman Coulter)-RNA. RNA library was eluted and loaded onto a flow cell for sequencing. Sequencing of the RNA was performed on a single R9.4/FLO-MIN106 flow cell on a MinION Mk1B for ~8 hours.

### Bioinformatics and real-time analysis

Immediately after loading the library onto a flow cell and sequencing, base-calling was started using the local-based software, Albacore version 2.3.1 (ONT) with the (--disable_filtering −q 0) command modifications from default. We first filtered out the sequenced reads below a size of 200 bases to avoid short, low-quality reads. The remaining reads were mapped to viral genomes and the host genomes using Minimap2 version 2.9 (Li, 2018) with the −k 7 from virus and −k 14 for host command modifications from default. The aligned reads were processed in downstream analysis. The number of mapped reads in virus and hosts were measured by Samtools version 1.5 (Li et al., 2009). The read coverage was calculated using genomecov package in BEDtools version 2.27.1 (Quinlan and Hall, 2010) and visualized using IGV (Robinson et al., 2011). The reference-based assembly was performed using four rounds consensus polishing of the reference genomes by Racon version 1.2.1 (Vaser et al., 2017).

### Disclosure of potential conflicts of interest

No potential conflicts of interest were disclosed.

## FUNDING

This work was funded in part by the Helen Adams & Arkansas Research Alliance Endowed Chair, the National Institute of General Medical Sciences of the National Institutes of Health award no. P20GM125503 and National Institutes of Health award no R01AI103053.

